# Specificity and effector functions of non-neutralizing gB-specific monoclonal antibodies isolated from healthy individuals with human cytomegalovirus infection

**DOI:** 10.1101/2020.01.14.907204

**Authors:** Matthew L. Goodwin, Helen S. Webster, Hsuan-Yuan Wang, Jennifer A. Jenks, Cody S. Nelson, Joshua J. Tu, Jesse Mangold, Sarah Valencia, Jason S. McLellan, Daniel Wrapp, Tong-Ming Fu, Ningyan Zhang, Daniel C Freed, Dai Wang, Zhiqiang An, Sallie R. Permar

**Affiliations:** Duke Human Vaccine Institute, Duke University Medical Center, Durham, NC, USA; Department of Molecular Biosciences, The University of Texas at Austin, Austin, TX, USA; Merck & Co., Inc., Kenilworth, NJ, USA; Texas Therapeutics Institute, Brown Foundation Institute of Molecular Medicine, The University of Texas Health Science Center, Houston, USA

## Abstract

Human cytomegalovirus (HCMV) is the most common congenital infection, and the leading nongenetic cause of sensorineural hearing loss (SNHL) in newborns globally. A gB subunit vaccine administered with adjuvent MF59 (gB/MF59) is the most efficacious tested to-date, achieving 50% efficacy in preventing infection of HCMV-seronegative mothers. We recently discovered that gB/MF59 vaccination elicited primarily non-neutralizing antibody responses, that HCMV strains acquired by vaccinees more often included strains with gB genotypes that are distinct from the vaccine antigen, and that protection against HCMV acquisition correlated with ability of vaccine-elicited antibodies to bind to membrane associated gB. Thus, we hypothesized that gB-specific non-neutralizing antibody binding breadth and function are dependent on their epitope and genotype specificity as well as their ability to interact with membrane-associated gB. Twenty-four gB-specific monoclonal antibodies (mAbs) isolated from naturally HCMV-infected individuals were mapped for gB domain specificity by binding antibody multiplex assay (BAMA) and for genotype preference binding to membrane-associated gB presented on transfected cells. We defined their non-neutralizing functions including antibody dependent cellular phagocytosis (ADCP) and antibody dependent cellular cytotoxicity (ADCC). The isolated gB-specific non-neutralizing mAbs were primarily specific for Domain II and linear antigenic domain 2 site 2 (AD2). We observed variability in mAb gB genotype binding preference, with increased binding to gB genotypes 2 and 4. Functional studies identified two gB-specific mAbs that facilitate ADCP and have binding specificities of AD2 and Domain II. This investigation provides novel understanding on the impact of gB domain specificity and antigenic variability on gB-specific non-neutralizing antibody responses.

**Importance:** HCMV is the most common congenital infection worldwide, but development of a successful vaccine remains elusive. gB-specific non-neutralizing mAbs, represent a distinct anti-HCMV Ab subset implicated in the protection against primary infection during numerous phase-II gB/MF59 vaccine trials. By studying non-neutralizing gB-specific mAbs from naturally infected individuals, this study provides novel characterization of binding site specificity, genotypic preference, and effector cell functions mediated by mAbs elicited in natural infection. We found that a panel of twenty-four gB-specific non-neutralizing mAbs bind across multiple regions of the gB protein, traditionally through to be targeted by neutralizing mAbs only, and bind differently to gB depending if the protein is soluble versus embedded in a membrane. This investigation provides novel insight into the gB-specific binding characteristics and effector cell functions mediated by non-neutralizing gB-specific mAbs elicited through natural infection, providing new endpoints for future vaccine development.

## Introduction

Each year, an estimated 40,000 children in the U.S. are born with congenital human cytomegalovirus (HCMV) infection (cCMV), with roughly 8,000 afflicted children developing long term sequelae of disease such as sensorineural hearing loss, neurodevelopmental delay, visual impairment, and psychomotor disability [1, 2]. While mothers with primary HCMV infection during pregnancy have rates of vertical transmission up to 40%, mothers with HCMV reactivation or reinfection can also transmit the virus to their developing infant but at a rate in the range of <1-4% [3–5]. Additionally, HCMV infection post-transplantation and consequently graft rejection, remain significant complications for solid-organ transplant patients, especially for those HCMV naïve recipients with transplants from HCMV seropositive donors [6, 7]. Although the protective correlates of immunity against HCMV infection have not been fully elucidated, there is ample evidence that immune factors elicited by natural infection can confer some degree of protection for mothers, infants, and immunosuppressed individuals against HCMV reinfection or reactivation. Thus, understanding how natural immunity against HCMV mediates protection may offer the basis for development of an effective HCMV vaccine.

Numerous HCMV vaccine candidates have been tested clinically, including live-attenuated virus, viral glycoprotein subunit formulations, viral vectors, and single/bivalent DNA plasmids, yet few have demonstrated sufficient protection as compared to natural immunity to warrant late stage clinical development [8]. Interestingly, the glycoprotein B (gB/MF59) subunit vaccine achieved nearly 50% efficacy in preventing HCMV primary infection in distinct cohorts of HCMV seronegative postpartum women and adolescent girls [9, 10]. Moreover, in HCMV seronegative patients receiving organs from HCMV seropositive donors, the gB/MF59 vaccine was shown to reduce the magnitude and duration of posttransplant HCMV viremia [11]. gB is a viral fusion protein, essential for viral infection of host cells, and is a target of both neutralizing and non-neutralizing antibodies [12]. Counterintuitively, the gB/MF59 vaccination elicited limited neutralizing responses when compared with HCMV seropositive individuals [13, 14]; yet, gB/MF59 vaccination elicited effector-cell mediated antibody responses, including antibody dependent cellular phagocytosis (ADCP). This observation prompted a hypothesis that non-neutralizing antibodies against gB may contribute to protection against HCMV acquisition [13, 14]. Moreover, effector antibody functions mediated by NK cells may be crucial for targeting HCMV-infected cells, further underlying the significance of non-neutralizing antibodies in anti-HCMV immunity [15, 16].

gB is a homotrimeric class III fusion protein that contains 5 defined antigenic domains, namely AD1 - AD5, with AD2 site 1, AD4 (Domain II), and AD5 (Domain I) as the target of neutralizing antibodies, and AD2 site 2, AD1, and the furin cleavage site as reported primary targets of non-neutralizing antibodies [12, 17]. The transmembrane and intravirion domains of gB (AD3), are thought to be obscured from antibody recognition in the membrane-associated portion. Yet, the epitope-specific antibody responses stimulated by natural HCMV infection and gB/MF59 vaccination demonstrate AD3 immunodominance, with 76% of total linear gB-specific antibody responses directed at this region in naturally HCMV infected vs 32% in gB/MF59 vaccinated individuals [12]. Variation in epitope-specific antibody binding may also stem from gB genotype-specific differences amongst clinical HCMV strains [17]. In fact, differences amongst HCMV gB genotypes correlate with cell tropism during HCMV infection [18]. Interestingly, it was recently reported that the ability of gB/MF59 vaccine sera to bind to membrane associated gB, via a transfected cell binding assay, predicts risk of HCMV acquisition [19]. Furthermore, like HSV gB and VSV protein G, HCMV gB is undergoes significant conformational transition from prefusion to postfusion states to mediate viral and host-cell membrane fusion [20, 21]. Thus, the CMV vaccine field needs characterization of antibodies that bind distinctly to soluble postfusion gB constructs and cell-associated gB, with the latter possibility representing a distinct prefusion-like structure.

Because there are stark differences in the gB epitope binding profiles of naturally-infected individuals and gB/MF59 vaccinees [13, 22], closing the gap in our understanding of immune-protection offered by prior exposure to natural HCMV infection and vaccination will require investigation into how gB as an antigen induces potentially protective effector antibody responses. The aim of this study is to characterize the non-neutralizing gB-specific antibody responses in natural HCMV infection for a panel of gB-specific mAbs isolated by memory B cell cultures from three naturally infected individuals [23]. Towards this goal, we analyzed a panel of 24 mAbs by defining their target specificity for gB domains as well as genotype specificity, and analyzed effector functions for these mAbs in ADCC and ADCP assays. Understanding gB diversity, and how gB-specific antibodies bind and mediate antiviral functions across genotypes to different antigenic domains, will be critical if we hope to optimize anti-gB antibody responses in future iterations of a HCMV vaccine.

## Methods

### gB-specific monoclonal antibodies (mAbs)

gB-specific monoclonal antibodies (mAbs) were isolated from healthy adult volunteers with previous natural HCMV infection. Subjects were consented for blood sampling in accordance with NIH guidelines [23]. B cell isolation and recombinant monoclonal antibody isolation and production using a human IgG1 backbone was performed as previously described [23]. Briefly, total RNA from isolated memory B cells, which screened positive for binding activity against gB, was converted to cDNA using a reverse transcription kit (Invitrogen), and IgG genes were identified by PCR primers. After cloning the variable regions of gB-specific mAbs from B cells from three donors the VH and VL sequences were cloned into a human IgG1 vector and recombinantly expressed in HEK293 cells in Zhiqiang An’s laboratory at the University of Texas Health Science Center at Houston.

### Postfusion gB ectodomain trimer and gB domain protein expression and purification

A pαH expression plasmid encoding an artificial signal sequence, residues 32-692 of HCMV gB from strain AD-169 with solubilized fusion loops, as described previously [12, 24], and C-terminal 6×His and TwinStrep tags was used to transiently transfect 293F cells. Secreted postfusion gB trimers were purified using Strep-Tactin resin (IBA Life Sciences) before being run over a Superose 6 size-exclusion column (GE Healthcare) in 2 mM Tris pH 8.0, 200 mM NaCl and 0.02% NaN_3_. Sequences encoding HCMV glycoprotein B (gB) Domain II (AA 112-133 + 143-438) from the Merlin strain was tagged at the 5’ end with the UL132 signal peptide sequence MPAPRGLLRATFLVLVAFGLLLHMDFS and hemagglutinin (HA) tag, and at the 3’ end with an avidin and polyhistidine tag. The discontinuous sequence encoding gB Domain II (AA 112-133 + 343-438) was joined with the flexible linker Ile-Ala-Gly-Ser-Gly. For Merlin strain gB Domain I (AA133-343), the 5’ HA and 3’ avidin/polyhistidine tags omitted due to hypothesized steric hinderance. Nucleotides were codon optimized for mammalian cells, synthesized *de novo* (Genscript), then cloned into pcDNA3.1(+) mammalian expression vector (Invitrogen) via *Bam*HI at the 5′ end and *Eco*RI site at the 3′ end. Plasmids were transiently transfected into 293F suspension cells as previously described using polyethyleneimine transfection reagent (Sigma-Aldrich) [25]. Supernatant was harvested 5 days later, and purified using Nickel-NTA resin for gB Domain II (Thermo Fisher Scientific), and lectin resin (VWR) for gB Domain I. Purity and identity were confirmed by Western blot using polyclonal CMV IgG (Cytogam – CSL Behring) or monoclonal antibodies SM10 (Domain I) and SM5-1 (Domain II).

### Mapping gB-specific mAb domain specificity by BAMA

Antibody responses against full length gB protein (only missing the transmembrane domain), gB ectodomain, and gB domains were measured as previously described [26]. Carboxylated fluorescent beads (Luminex) were covalently coupled to purified HCMV antigens and incubated with monoclonal antibodies in assay diluent (phosphate-buffered saline, 5% normal goat serum, 0.05% Tween 20, and 1% Blotto milk). The gB domain antigen panel, included full length gB (generously provided by Sanofi), gB ectodomain, gB domain I, gB domain II, gB AD-1 (myBiosource), and biotinylated linear gB AD-2 site 2 (biotin-AHSRSGSVQRVTSS), and biotinylated linear gB AD-2 site 1 (biotin-NETIYNTTLKYGD) are previously described [13]. All gB proteins are based on the Towne strain gB (genotype 1). CMV glycoprotein–specific antibody binding was detected with phycoerythrin-conjugated goat anti-human IgG (2 μg/mL, Southern Biotech). Beads were washed and acquired on a Bio-Plex 200 instrument (Bio-Rad), and results were expressed as mean fluorescence intensity. A panel CMV seronegative plasma samples (n=30) were included to determine nonspecific baseline levels of binding. Minimal background activity was observed, so the threshold for positivity for each antigen was set at the mean value (100 MFI) of negative control sera to each antigen + two standard deviations. Blank beads were used in all assays to account for nonspecific binding. All assays included tracking of CMV immunoglobulin (Cytogam, generously gifted by CSL Behring) standard by Levy-Jennings charts. The preset assay criteria for sample reporting were coefficient of variation per duplicate values of ≤20% for each sample and ≥100 beads counted per sample. All mAbs were analyzed at a concentration of 30 μg/mL for each antigen: full-length gB (gift of Sanofi), gB ectodomain, gB Domain I, gB Domain II, gB AD-1 and gB AD-2 sites 1 and 2. This concentration was predetermined to be within the linear range of binding based on testing serial dilutions of a small subset of gB specific mAbs (1-155, 1-235, 2-43, 1-189, 3-54, and 3-74).

### gB-specific mAb binding strength measured via ELISA

gB-specific monoclonal antibody binding responses were measured against full length gB protein (Sanofi), gB regional epitopes (Ectodomain, Domain I), or gB peptides (AD2 site 1, AD2 site 2). All gB constructs were solubilized in 0.1 M NaHCO_3_, with proteins plated at a concentration of 3 μg/mL and peptides at 10 μg/mL respectively. Plates were washed then blocked before adding gB-specific mAbs at either 5 μg/mL for conformational protein ELISAs or 100 μg/mL for peptide ELISAs. After incubation with mAbs, plates were washed and incubated with a 1:5000 goat-anti human HRP conjugated IgG secondary (Jackson ImmunoResearch). ELISAs were developed with SeraCare ELISA kit and read at 450 nm.

### Assessment of gB-specific mAb avidity via surface plasmon resonance (SPR)

Postfusion trimeric gB ectodomain was captured on an NTA sensor chip to ~500 response units (RUs) per cycle using a Biacore X100 (GE Healthcare). The chip was doubly regenerated using 0.35 M EDTA and 0.1 M NaOH followed by 0.5 mM NiCl_2_. Three samples containing only buffer were injected over both ligand and reference flow cells, followed by single injections of each mAb at a concentration of 25 nM. Samples that did not initially result in interpretable sensorgrams were repeated using a concentration of 250 nM. The resulting data were double-reference subtracted and fit to a 1:1 binding model using the Biacore X100 Evaluation software.

### gB genotype-specific transfected cell binding

HEK293T cells were cultured overnight to ~50% confluency in a T25 flask, and then co-transfected using TransIT-mRNA Transfection Kit (Mirus Bio) with a GFP-expressing mRNA plasmid (Miltenyi Biotec) and a second plasmid encoding either full length gB from genotypes 1, 2, 3, 4, or 5 (UPenn, Drew Weissman). Transfected cells were incubated at 37°C and 5% CO_2_ for 24 hours, washed with PBS once, and detached using 0.05% trypsin + EDTA (Thermo Fisher Scientific). Cells were re-suspended in DMEM complete medium and counted using Countess Automated Cell Counter (Invitrogen). 100,000 cells were placed in tubes and stained with LIVE/DEAD Aqua Dead Cell Staining Kit (Thermo Fisher Scientific) diluted 1:1000 at room temperature for 20 minutes. 50,000 live cells were placed in each well of a 96-well V-bottom plate (Corning). The plates were centrifuged at 1,200 x g for 5 minutes and the supernatants were aspirated. Cells were incubated with monoclonal antibodies, which were diluted to 5 μg/ml in duplicate in DMEM complete medium, at 37°C and 5% CO_2_ for 2 hours. After washing with wash buffer (PBS + 1% FBS) twice, cells were incubated with PE-conjugated mouse anti-human IgG Fc (Southern Biotech) diluted 1:200 at 4°C for 30 minutes. Following two additional wash steps, cells in tubes and plates were resuspended and fixed in PBS + 10% formalin for 10 minutes at room temperature. Fixed cells were washed once and resuspended in wash buffer for flow cytometry. Events were acquired on LSR II machine (BD biosciences) using high-throughput sampler (HTS). Data were analyzed with Flowjo software (Tree Star, Inc.), and the PE+ population was identified from the live GFP+ cell population for each sample. Non-specific binding of PE-conjugated mouse anti-human IgG Fc was corrected in the analysis.

### Natural killer (NK) cell CD107a degranulation assay

Cell-surface expression of CD107a was used as a marker for NK cell degranulation (7, 8). ARPE cells were plated at 3×10_4_ cells/well in a 96-well flat-bottom tissue culture plate and allowed to incubate for 24 hours at 37°C. Cells were infected with AD169r-GFP at an MOI of 1.0 or transfected with gB-encoding mRNA using the TransIT mRNA transfection kit (Mirus Biosciencies), then incubated a further 48 hours at 37°C. Following incubation, supernatant was removed and the transfected or infected cell monolayers were washed once with RPMI 1640 containing 10% FBS, HEPES, Pen-Strep-L-Glut, Gentamicin (R10 media) before addition of NK cells. Primary human NK cells were isolated from blood peripheral mononuclear cells (PBMC) after overnight rest in R10 media with 10ng/mL IL-15 (Miltenyi Biotech) by depletion of magnetically labeled cells (Human NK cell isolation kit, Miltenyi Biotech). 5×10_4_ live NK cells were added to each well containing gB transfected or HCMV-infected ARPE19 cell monolayers. mAbs were diluted in R10 and added to the cells at a final dilution of 25μg/mL in duplicate. Brefeldin A (GolgiPlug, 1 μl/ml, BD Biosciences), monensin (GolgiStop, 4μl/6mL, BD Biosciences), and CD107a-FITC (BD Biosciences, clone H4A3) were added to each well and the plates were incubated for 6 hours at 37°C in a humidified 5% CO_2_ incubator. NK cells were then gently resuspended, taking care not to disturb the ARPE-19 cell monolayer, and the NK containing supernatant was collected and transferred to 96-well V-bottom plates. The recovered NK cells were washed with PBS, and stained with LIVE/DEAD Aqua Dead Cell Stain at a 1:1000 dilution for 20 minutes at room temperature. The cells were then washed with 1%FBS PBS and stained for 20 minutes at room temperature with the following panel of fluorescently conjugated antibodies diluted in 1%FBS PBS: CD56-PECy7 (BD Biosciences, clone NCAM16.2), CD16-PacBlue (BD Biosciences, clone 3G8), and CD69-BV785 (BioLegend, Clone FN50). The cells were then washed with 1%FBS PBS, fixed and permeabilized for 20 minutes at 4°C (BD Fixation/Permeabilization solution), washed with BD Perm/Wash, and stained for intracellular interferon (IFN)-γ-BV711 (BioLegend, clone 4S.B3) and tumor necrosis factor-alpha (TNF)-α-BV650 (BD Biosciences, clone Mab11) for 30 minutes at 4°C. The cells were then washed twice and re-suspended in 1% paraformaldehyde fixative for flow cytometric analysis. Data analysis was performed using FlowJo software (v9.9.6). Data is reported as the % of CD107a+ live NK cells (singlets, lymphocytes, aqua blue-, CD56+ and/or CD16+, CD107a+). The threshold for positivity is the mean response of preimmune samples plus two standard deviations. CD69, TNFα, and IFNγ were not included in the final analysis due to the low frequency of CD107a+ responses.

### Antibody dependent cellular phagocytosis (ADCP)

Approximately 3.5×10_6_ PFU of concentrated, sucrose gradient-purified AD169r-GFP virus was transferred to a 100,000 kDa Amicon filter (Millipore), then buffer exchanged with 1× PBS, concentrated down to approximately 100 μL, and transferred to a microcentrifuge tube. Next, 10 μg of AF647 NHS ester (Invitrogen) reconstituted in DMSO was added to the concentrated, purified virus for direct fluorescent conjugation, then this reaction mixture was incubated at room temperature for 1 hour with constant agitation. The reaction was quenched with 80 μL of 1 M Tris-HCl, pH 8.0, then the fluorophore-labelled virus was diluted 25x in wash buffer (PBS + 0.1% FBS). mAbs were diluted to 0.1 mg/mL in wash buffer, then 10 μL of each diluted mAb was combined with 10 μL of diluted, fluorophore-conjugated virus in a round-bottom, 96-well plate and allowed to incubate at 37°C for 2 hours. Following this incubation step, 50,000 THP-1 cells were added to each well, suspended in 200 μL primary growth media. Plates were centrifuged at 1200x g and 4°C for 1 hour in a spinoculation step, then incubated at 37°C for an additional hour. Cells were re-suspended and transferred to a 96-well V-bottom plate, then washed twice prior to fixation in 100 μL DPBS + 1% formalin. Events were acquired on LSR II machine (BD biosciences) using the HTS. The % AF647+ cells was calculated from the full THP-1 cell population and reported for each sample. A cutoff for a sample mediating ADCP was defined as > 99% AF647+ signal from THP1 cells incubated with fluorophore conjugated virus and an HCMV seronegative control.

## Results

### Mapping the domain-specificity of non-neutralizing gB-specific mAbs from HCMV seropositive individuals

We first mapped the binding specificity of the panel of gB-specific mAbs for defined neutralizing and non-neutralizing antigenic domains of gB by BAMA. The most frequent binding specificity was against Domain II (37.5%), as well as AD2 site 2 (12.5%) and AD3/MPER (12.5%) (Figure 1A). While 4 mAbs demonstrated detectable binding to both linear AD2 Site 1 and Site 2, these mAbs are likely AD2 Site 2 specific, given that previously described AD2 Site 1 mAbs have neutralizing activity [27, 28]. Interestingly, there were three mAbs, 1-155, 1-237, and 3-18, which bound full length gB, (Sanofi), but did not bind a gB ectodomain that excludes the membrane proximal region (MPER) and the cytodomain containing AD3, suggesting specificity for one of these regions.

**Figure 1.**
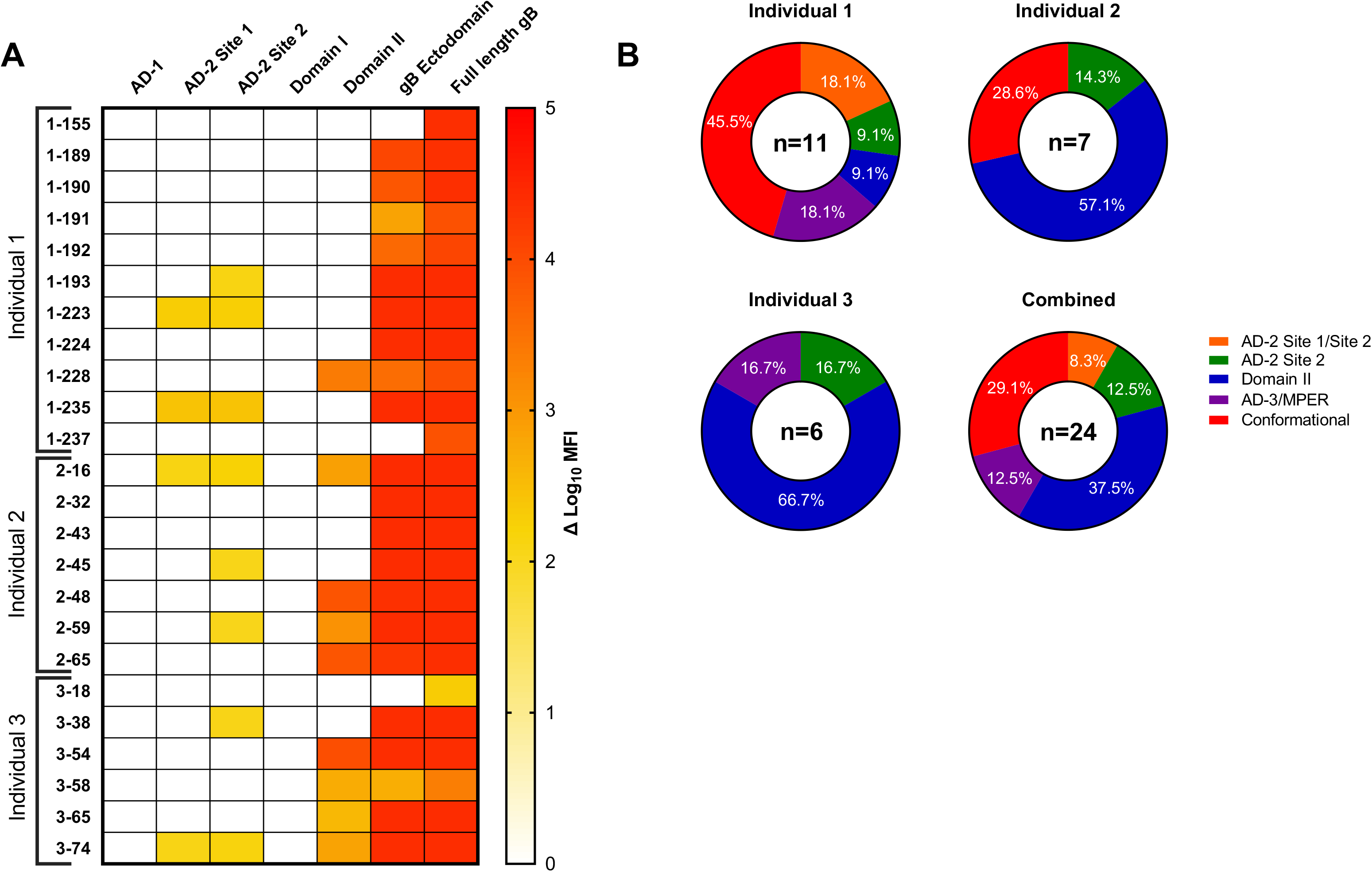
gB domain-specific mAb binding determined by BAMA. (A) Heat map of gB domain-specific binding strength for gB-specific mAbs represented as log_10_ mean fluorescent intensity (MFI). (B) Pie charts representing the percentage of total gB-specific mAbs binding to each domain specificity assessed by BAMA per each naturally HCMV infected individual and all individuals combined. AD-2 Site 1/Site 2: Binding to both the AD-2 Site 1 and Site 2 peptide, AD-2 site 2: binding to the site 2 peptide only, Domain II: binding to Domain II only, AD3/MPER: binding to full length gB, but not gB ectodomain, Conformational: binding to full length gB and gB ectodomain, but no other domain.

There are two groups of clonal populations of mAbs found in Individual 1, including mAbs 1-189, 1-190, 1-191, 1-192 which bind full length gB and gB ectodomain conformationally, but not to a defined epitope (termed “conformational” binding), and mAbs 1-223 and 1-224. Antigenic domain-specificity of gB-specific mAbs was heterogenous in Individual 1, but was dominated by Domain II in individuals 2 (57.1%) and 3 (66.7%) (Figure 1B). The binding profile for the combined panel of 24 gB-specific mAbs notably excludes any AD1 or AD5 (Domain I) binding mAbs, despite the polyclonal CMV-IgG preparation Cytogam binding to all antigens (data not shown).

### mAb binding kinetics to full length gB and trimeric post-fusion gB ectodomain

Next, we sought to describe the binding kinetics of each gB-specific mAb to the full length gB protein vs a trimeric post-fusion gB ectodomain construct. By understanding discrepancies in binding between full length gB and the post-fusion gB ectodomain, we aimed to highlight novel HCMV AD3 or MPER-specific mAbs and their binding kinetics. Multiple mAbs (1-155, 1-189, 1-191, 1-192, 1-237) exhibited poor binding strength to post-fusion gB (ectodomain), but retained robust binding to full length gB. MAbs 3-18 and 3-58 exhibited poor gB binding to both full length gB and post fusion gB ectodomain (Table 1).

**Table 1.** gB-specific mAb immunogenetics, binding strength, and avidity to full length gB and gB ectodomain a. Heavy chain and light chain genes, neutralization IC50 (μg/mL), as well as CDRH3 length listed as described in (Xia et al 2017). b. binding to full length gB and post fusion gB ectodomain EC_50_ (μg/mL) measured by ELISA. c. binding kinetics to post fusion gB ectodomain measured by SPR

| Monoclonal Antibody | Heavy Chain <sup>a</sup> | Light Chain <sup>a</sup> | CDRH3 Length <sup>a</sup> | Neutralization IC<sub>50</sub> (μg/mL) <sup>a</sup> | Epitope Specificity | Full length gB binding EC<sub>50</sub> (μg/mL) <sup>b</sup> | gB ectodomain binding EC<sub>50</sub> (μg/mL) <sup>b</sup> | gB ectodomain apparent avidity KD (nM) <sup>c</sup> |
| --- | --- | --- | --- | --- | --- | --- | --- | --- |
| 1-155 | IGHV1-69*01 | IGLV2-11*01 | 14 | 0 | AD3/MPER | 0.002 | 6.903 | 6.2 |
| 1-189 | IGHV1-69*01 | IGKV4-1*01 | 11 | 0 | Conformational | 0.007 | 2.562 | 702 |
| 1-190 | IGHV1-69*01 | IGKV4-1*01 | 11 | 0 | Conformational | 0.019 | 3.716 | 1.88 |
| 1-191 | IGHV1-69*01 | IGKV4-1*01 | 11 | 0 | Conformational | 0.009 | 6.163 | 10.8 |
| 1-192 | IGHV1-69*06 | IGKV4-1*01 | 11 | 0 | Conformational | 0.010 | > 100 | 116 |
| 1-193 | IGHV1-18*01 | IGKV2-30*01 | 22 | 1 | AD2 Site 2 | 0.003 | 0.004 | 0.0003 |
| 1-223 | IGHV1-69*03 | IGLV2-23*01 | 22 | 8.3 | AD2 Site 1/Site 2 | 0.009 | 0.013 | 0.301 |
| 1-224 | IGHV1-69*03 | IGKV3-20*01 | 22 | 6.4 | Conformational | 0.006 | 0.011 | 0.116 |
| 1-228 | IGHV3-13*01 | IGKV1-33*01 | 10 | 0 | Domain II | 0.001 | 8.014 | N.D. |
| 1-235 | IGHV4-31*03 | IGKV4-1*01 | 11 | 0 | AD2 Site 1/Site 2 | 0.001 | 0.002 | 0.019 |
| 1-237 | IGHV1-69*01 | IGKV4-1*01 | 12 | 0 | AD3/MPER | 0.003 | 2.656 | 5.27 |
| 2-16 | IGHV1-69*01 | IGKV3-15*01 | 15 | 11 | Domain II | 0.002 | 0.004 | 0.0004 |
| 2-32 | IGHV4-59*01 | IGKV4-1*01 | 12 | 0 | Conformational | 0.003 | 0.006 | N.D. |
| 2-43 | IGHV6-1*01 | IGKV4-1*01 | 14 | 15 | Conformational | 0.003 | 0.004 | 0.0034 |
| 2-45 | IGHV4-34*02 | IGKV3-15*01 | 24 | 9.8 | AD2 Site 2 | 0.003 | 0.006 | 0.0004 |
| 2-48 | IGHV1-2*02 | IGKV3-20*01 | 14 | 14.6 | Domain II | 0.003 | 0.006 | 0.113 |
| 2-59 | IGHV4-34*02 | IGLV1-47*01 | 18 | 2.3 | Domain II | 0.006 | 0.010 | 0.0016 |
| 2-65 | IGHV1-2*02 | IGKV3-20*01 | 14 | 6.4 | Domain II | 0.007 | 0.018 | 0.081 |
| 3-18 | IGHV3-21*01 | IGKV1D-8*01 | 26 | 0 | AD3/MPER | 16.370 | > 100 | 22.5 |
| 3-38 | IGHV1-18*01 | IGKV4-1*01 | 14 | 0 | AD2 Site 2 | 0.006 | 0.008 | 0.0003 |
| 3-54 | IGHV1-46*01 | IGKV1-5*03 | 22 | 1 | Domain II | 0.006 | 0.011 | 0.474 |
| 3-58 | IGHV3-30*03 | IGLV3-10*01 | 34 | 0 | Domain II | 5.733 | 20.560 | 1.82 |
| 3-65 | IGHV3-33*01 | IGKV4-1*01 | 16 | 0 | Domain II | 0.003 | 0.003 | 0.036 |
| 3-74 | IGHV3-30-3*01 | IGLV3-25*03 | 28 | 0 | Domain II | 0.601 | 0.723 | 1.52 |

Three isolated gB-specific mAbs with binding to both the linear AD2 Site 1 and Site 2 regions, regardless of individual donor, demonstrated both high binding strength and avidity for full length gB (EC_50_ 0.001 − 0.009 μg/mL) and gB ectodomain (0.002 − 0.013 μg/mL) (Table 1). gB mAbs with AD2 site 2 only specificity also had robust binding to full length gB (EC_50_ 0.003 – 0.006 μg/mL) as well as gB ectodomain (0.004 – 0.008 μg/mL) (Table 1). Of the mAbs which bound Domain II, all except three, 1-228, 3-58 and 3-74, showed strong binding to both full length gB and gB ectodomain (EC_50_ < 0.020 μg/mL) (Table 1). Specifically, mAb 3-74, which binds Domain II predominantly, demonstrated poor binding and avidity to both full length gB and gB ectodomain by ELISA and SPR. Two Domain II-specific mAbs bound with an interesting pattern to gB ectodomain. Domain II-specific mAb 1-228 had strong binding to full length gB (EC_50_ 0.001 μg/mL) but very poor binding to gB ectodomain by ELISA (EC_50_ 8.014 μg/mL) and no detectable (N.D.) avidity by SPR (Table 1). Further, Domain II-specific mAb 3-58 had poor binding strength to both full length gB (EC_50_ 5.73 μg/mL) and gB ectodomain (EC_50_ 20.56 μg/mL) (Table 1).

MAbs which bound to the full length gB and not the ectodomain in the gB domain mapping, and therefore potentially bind AD3/MPER, 1-155 and 1-237, followed predictable patterns of binding with strong binding to full length gB (EC_50_ < 0.004 μg/mL) and negligible binding strength and avidity to gB ectodomain (Figure 1A, Table 1). mAb 3-18, also determined to be AD3/MPER specific by BAMA, demonstrated poor binding measured by both ELISA (EC_50_ > 100 μg/mL) and SPR (*K*_D_ 22.5 nM) to the ectodomain as well Finally, those mAbs which didn’t have a definable epitope specificity by gB domain mapping, and termed “conformational”, had highly variable binding strength to gB ectodomain, but fairly strong binding to full length gB. Interestingly, mAb 2-32, which bound both full length gB (EC_50_ 0.003 μg/mL) and gB ectodomain (EC_50_ 0.006 μg/mL) well via ELISA, as well as has binding detected to both the full length gB and gB ectodomain antigens via BAMA (Figure 1A), demonstrated undetectable binding to gB ectodomain by SPR (N.D., not detectable).

### Cell associated gB DNA transfected-cell binding

We have recently discovered that the magnitude of binding to gB DNA transfected cells is a correlate of protection against primary HCMV acquisition in postpartum and adolescent women vaccinated with gB/MF59 in phase II clinical trials [19]. Next, we investigated whether binding to cell associated gB is dependent upon gB-specific mAb antigenic site specificity. gB-specific mAbs with AD-2 specificity demonstrated high magnitude binding to cell associated gB more consistently than mAbs from any other specificity represented in our panel (Figure 2A). AD-3 specific mAbs bind poorly to cell associated gB, consistent with epitope location in the transmembrane and cytosolic compartment [12]. Notably, binding to cell associated gB was not correlated to strength of binding to soluble gB measured by ELISA (Figure 2B). These findings demonstrate that gB-mAbs of various specificities, except for AD3, are capable of binding cell associated gB with high magnitude.

**Figure 2.**
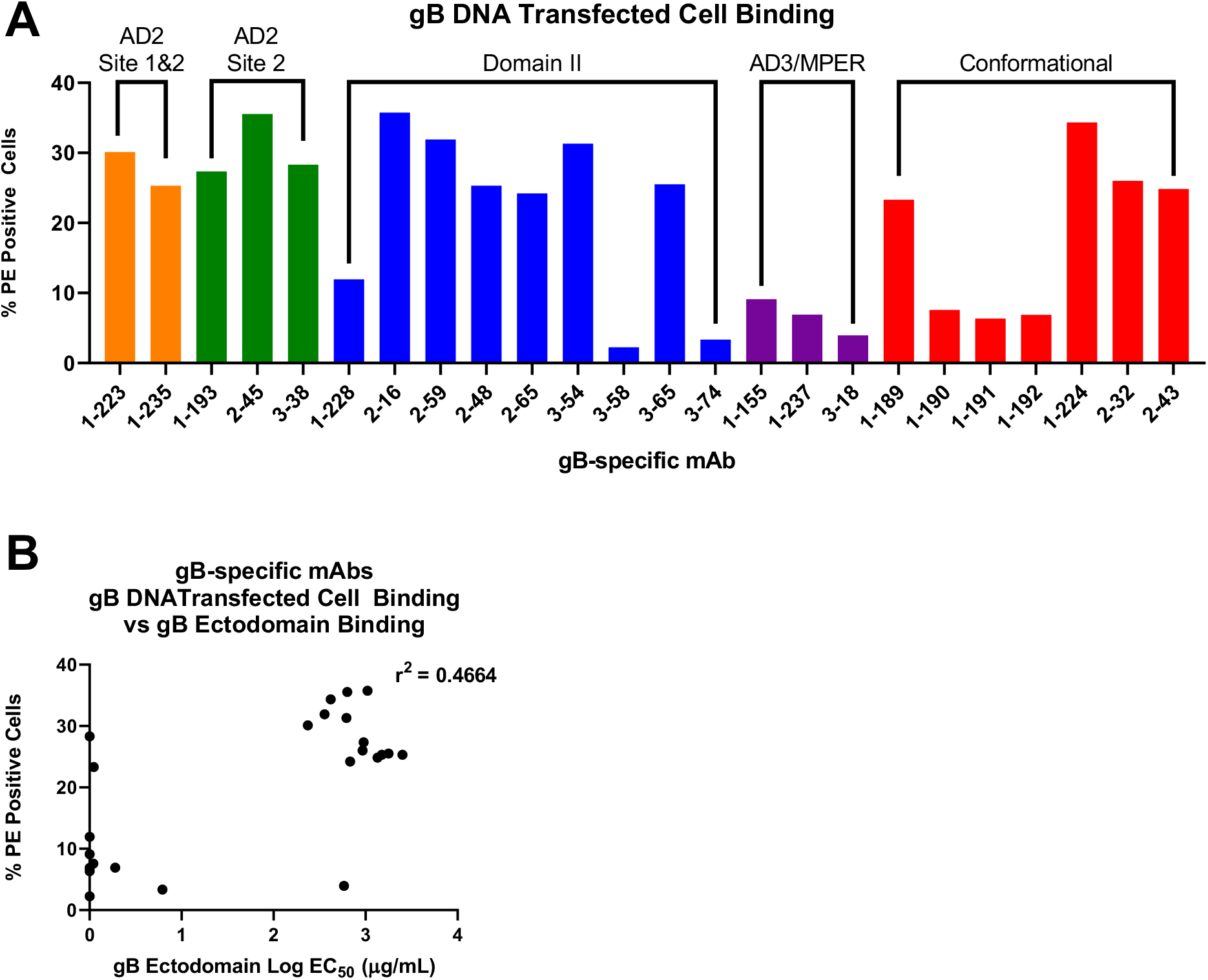
gB DNA transfected cell binding. (A) gB-specific mAb binding to cell-associated gB on the surface of gB DNA transfected cells. (B) gB-specific mAbs generally bind with high magnitude to cell-associated gB, but this binding strength is not highly correlated to strength of binding (EC_50_) to soluble gB ectodomain measured by ELISA.

### Cell associated gB genotype-specific mAb binding

We previously reported that gB/MF59 vaccinees may have had reduced acquisition of HCMV strains with gB1 genotype, the genotype matched to the vaccine construct, suggesting strain-specific protection [29]. As such, we next explored how non-neutralizing gB-specific mAbs differentially recognize gB genotypes 1-5, as expressed on the surface of a cell. Comparison of total, and genotype specific, gB transfected cell bound populations of each gB-specific mAb (Figure 3A) highlights significant variability for cell-associated gB binding across epitope specificities. While a majority of gB-specific mAbs bound all 5 genotypes, there was unequal binding magnitude across genotypes for the same mAb. To compare genotype preference amongst gB-specific mAbs (Figure 3B), we next assessed the percent representation of each genotype in the sum total gB genotype binding for each mAb. Here, there emerged a clear preference for gB-specific mAbs isolated from these naturally-infected donors for gB genotypes 2 and 4, representing the cumulative genotype preference for 11 of 14 gB-specific mAbs with gB genotype specific binding (Figure 3C). Genotype-dominant binding was determined by mAbs which bound a single gB genotype with a % PE positive population greater than 5x the % PE positive population of the lowest bound gB genotype. These findings are consistent across epitope specificities. Interestingly, mAbs which were unable to bind gB ectodomain measured by ELISA or BAMA, but bound full length gB and classified as AD3 specific, demonstrated equivalent cell associated gB binding across all tested gB genotypes (Figure 3A). These data implicate differences in epitope presentation of soluble gB and cell-associated gB.

**Figure 3.**
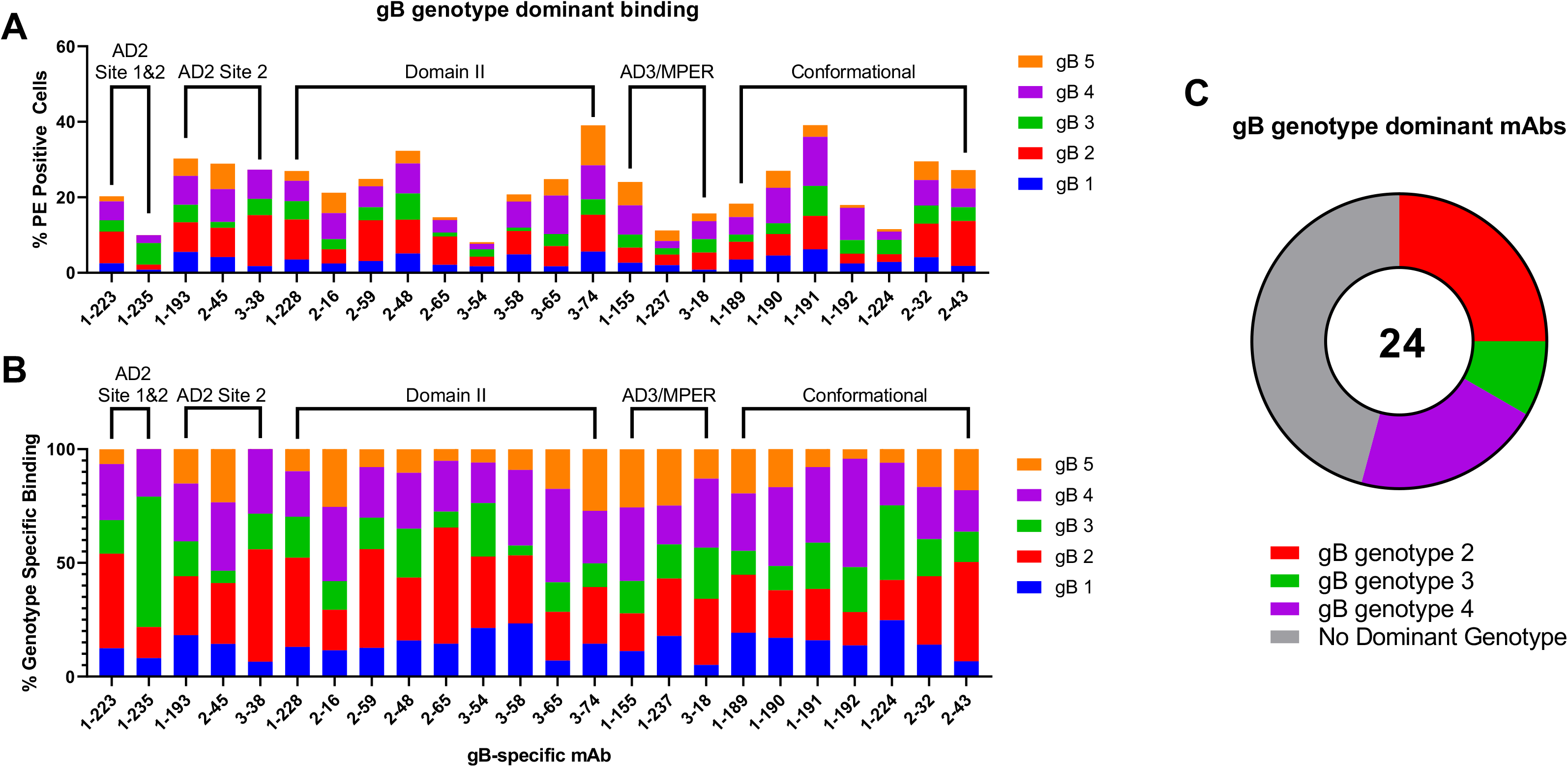
Cell associated gB genotype-specific mAb binding. (A) Percentage of gB genotype-specific transfected cells bound by gB-specific mAbs, grouped by domain specificity. (B) Percentage of gB genotype-specific transfected cells bound by the total population of gB transfected cells, demonstrating genotype preference for each gB-specific mAbs, listed by domain specificity. (C) gB-genotype preference for gB-specific mAbs defined if % PE positive population of a single transfected gB genotype was greater than 5 times the % PE positive population of the lowest bound cell-associated gB genotype by that mAb.

### Antibody dependent cellular cytotoxicity (ADCC)

Antibody-mediated NK cell cytotoxicity has been demonstrated as a crucial mechanism for controlling HCMV infection [30, 31]. To distinguish the domain specificity of gB-specific mAbs which mediate ADCC, we next screened our panel of mAbs for ability to mediate two well defined NK phenotypes of cytotoxic killing, CD107a upregulation, indicative of degranulation, (Figure 4A) and CD16 downregulation, indicative of NK cell activation (Figure 4B) in the presence of HCMV AD169r infected cell targets. While a number of mAbs across multiple domains demonstrated measurable CD107a degranulation, they were comparable to the negative control Synagis (mAb against respiratory syncytial virus). Next, a more specific marker of NK cell activation, CD16 downregulation calculated into a downregulation index (DRI), was measured for each gB-specific mAb. Notably, no gB-specific mAb demonstrated significant CD16 downregulation whereas HCMV hyperimmunoglobulin (Cytogam) had demonstrable activity.

**Figure 4.**
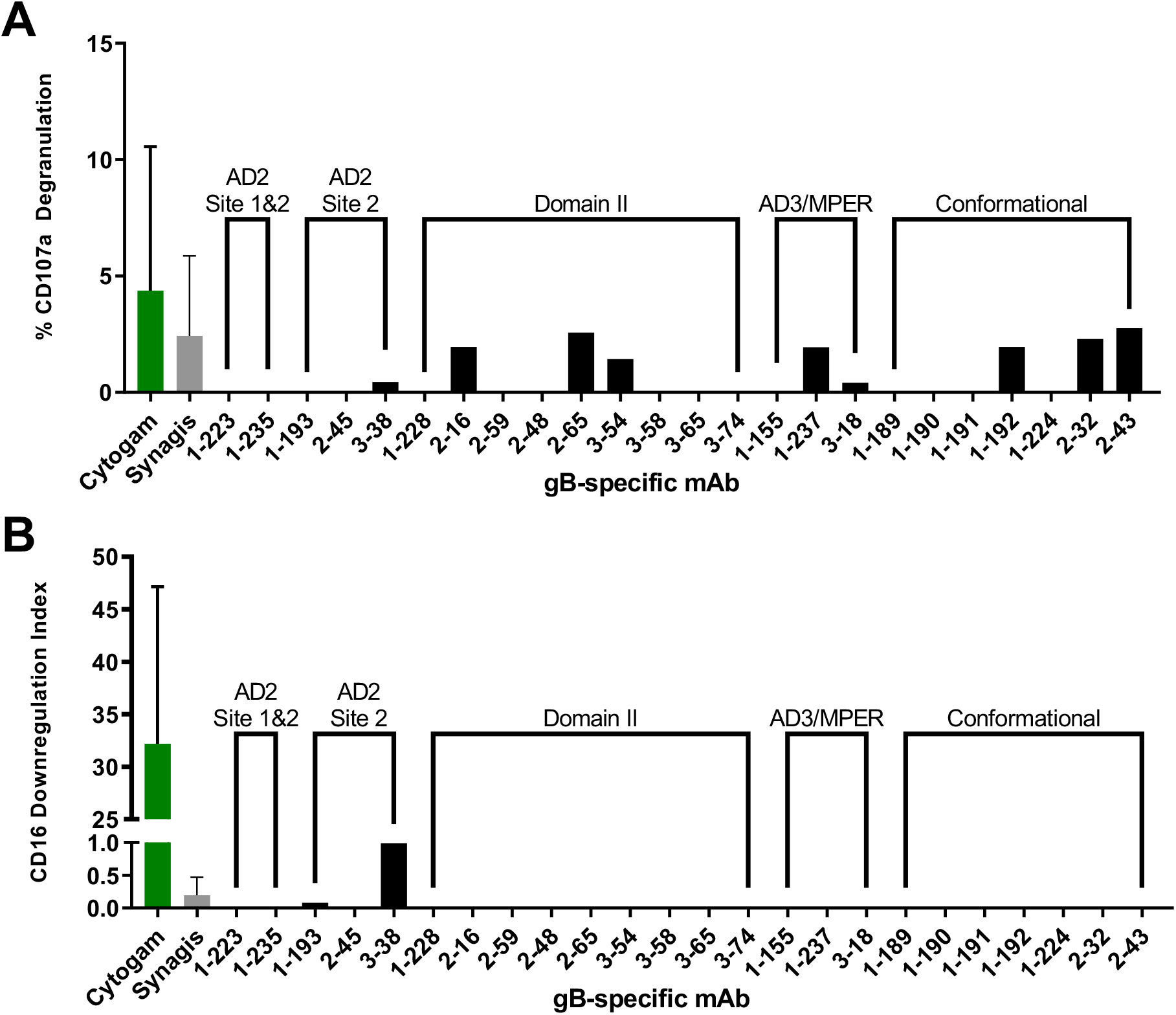
gB-specific mAbs mediate negligible NK activation ADCC responses when compared to polyclonal CMV-specific antibody preparation (Cytogam). gB-specific mAbs were screened for mediation of NK cell (A) CD107a degranulation and (B) CD16 downregulation (DRI) as measures of antibody-mediated NK cell activation. Data are presented as mAb specific response against AD169r-GFP infected ARPE19 cells minus mock infected negative control cells.

### gB AD2 Site 1/Site 2 and Domain II-specific mAbs 1-235 and 3-74 mediate ADCP

In a study of functional antibody responses to gB/MF59 vaccination, vaccinees were found to demonstrate limited neutralization responses, but robust ADCP [13]. To identify the domain specificity of gB-specific mAbs which can mediate ADCP, we screened our panel of gB-specific mAbs for ability to mediate whole virion phagocytosis (Figure 5A). Two gB-specific mAbs, 1-235 and 3-74, mediated phagocytosis of AD169r-GFP virions, which exceeded the baseline ADCP level of non-specific mAbs and seronegative control plasma by 19.4% and 14.7% respectively. To better assess their ADCP potency in both THP-1 cells and primary monocytes isolated from healthy donors (Figure 5B), mAbs 1-235 and 3-74 were titrated from a concentration of 100 μg/mL. ADCP increased in a dose-dependent manner for 1-235 and 3-74 in both THP1 cells and primary monocytes. Importantly, the gB epitope specificity of ADCP mediating mAb 1-235 was AD2 Site 1 and Site 2, while 3-74 specificity was Domain II (Figure 1A).

**Figure 5.**
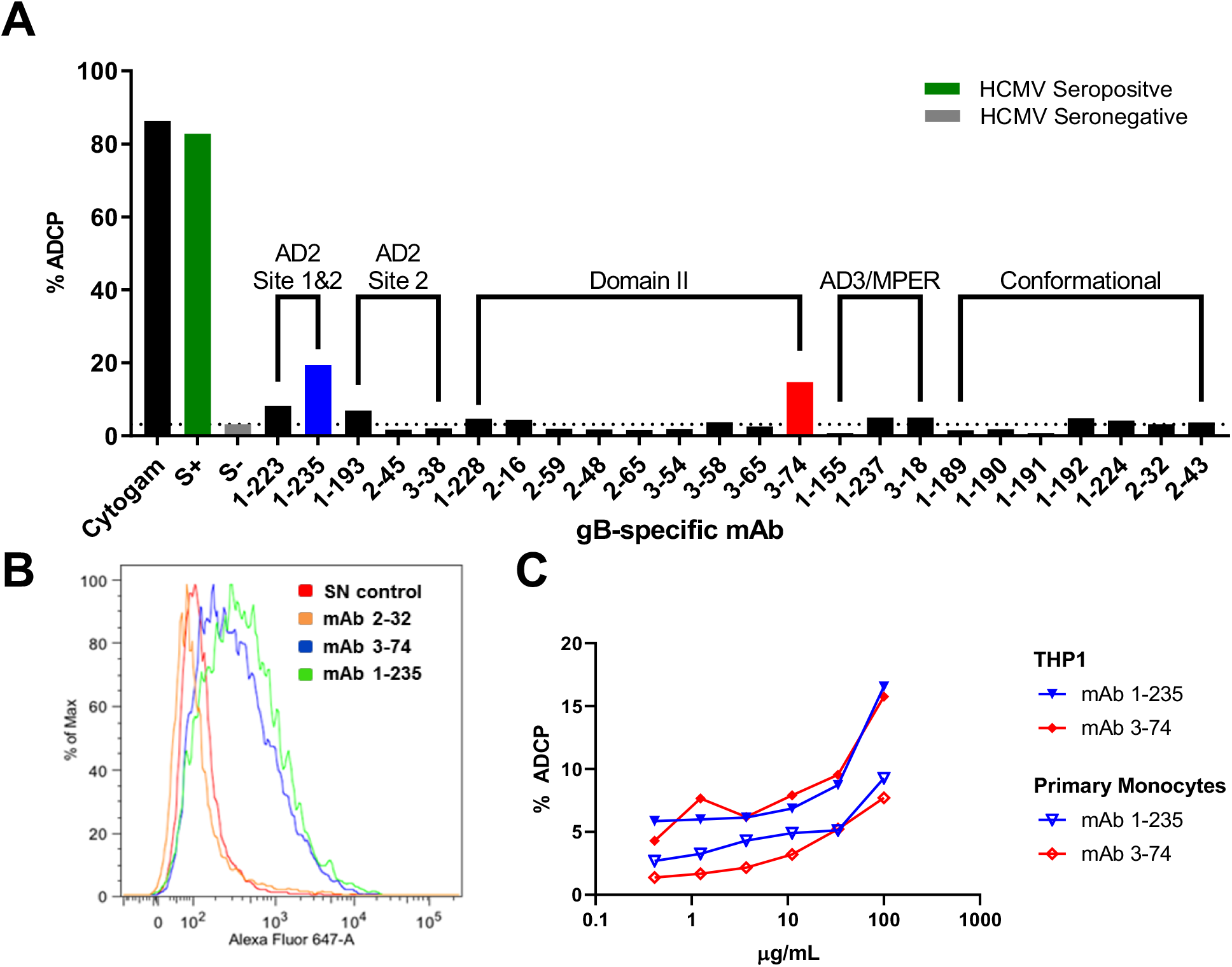
gB AD2 and Domain-1 specific mAbs 1-235 and 3-74 mediate antibody dependent cellular phagocytosis (ADCP). (A) gB-specific mAbs were screened for ADCP activity. (B) A representative histogram for ADCP mediating mAbs as compared to a HCMV seronegative control. The magnitude of ADCP was titrated over a dilution series for the two ADCP mediating mAbs using (C) THP1 cells (solid symbols) and primary monocytes (open symbols)

## Discussion

Identification of the dominant non-neutralizing antibody responses elicited by the partially-protective gB/MF59 vaccine has suggested a shift in the paradigm of immunogenicity endpoints for consideration when designing the next generation of HCMV vaccines [13]. While protective immunity through virus neutralization to prevent primary infection is considered the “gold standard” of HCMV vaccine development, next generation vaccine regimens will need to consider these non-neutralizing anti-viral antibodies. Indeed, strategies such as cell-to-cell spread enable immunologically-covert spread of infection even in the presence of neutralizing antibodies [32]. In fact, prophylactic passive immunization with both neutralizing and non-neutralizing mAbs were considered equally protective in a murine cytomegalovirus challenge model [33]. Accordingly, non-neutralizing antibodies which mediate effector functions like ADCP, like those described in this study, may have potential importance as immunological endpoints of a protective HCMV vaccine.

A majority (> 90%) of gB-specific antibodies from B cell clones have no neutralizing activity [34]. Indeed, the partially-protective gB/MF59 vaccine elicited limited heterologous neutralization [13]. While the specificity and characteristics of neutralizing gB-specific mAbs have been well described, the types of naturally-elicited gB-specific mAbs that mediate non-neutralizing effector functions which may be critical for protection against HCMV acquisition remain to be fully characterized.

In this study, 24 non-neutralizing gB-specific mAbs isolated from naturally HCMV-infected individuals were assessed for epitope binding specificity and affinity, gB genotype preference, and Fc-mediated effector functions. These non-neutralizing mAbs bound predominantly to Domain II, or to one or both of the binding sites of AD2. Domain II is largely the target of neutralizing antibodies, with one study identifying Domain II-specific binding by neutralizing antibodies in greater than 90% of HCMV seropositive subjects tested. Interestingly, three mAbs from the panel demonstrated robust binding to the full length gB protein but failed to bind a gB ectodomain, suggesting specificity within the AD3 or MPER region, a portion of the gB construct located in the cytodomain of membrane associated gB [12]. This potentially hidden region on an intact virion or infected cell could certainly be exposed through protein shedding from cell lysis or disruption of the HCMV virion [35], similarly to other viral structural antigens such as pp150 and pp28. However, it also raises questions regarding the structure of this membrane-associated region of the protein. Interestingly, none of the non-neutralizing gB-specific mAbs in this panel had appreciable binding to AD1 or Domain I. AD1 is targeted by both neutralizing and non-neutralizing mAbs [34, 36]. As such, the lack of Domain I and AD1 binding from this representative panel could be the result of sampling bias from only three individuals [23], by selection bias of excluding mAbs which bind to traditionally neutralizing domains, or non-optimal conformation of the protein used for B cell sorting to isolate these mAbs.

Antibodies mediating neutralization after gB/MF59 vaccination and natural infection can be strain specific [13, 37] and may not provide equivalent degrees of protection against all strains. It has been reported that breakthrough HCMV infections in the vaccine subjects receiving gB/MF59, which is a gB genotype 1 based vaccine, were more likely to be infected with HCMV strains expressing gB genotypes 3 or 5 than strains 1, 2, or 4 [29]; evidence suggesting gB/MF59 induced antibodies were limited due to its strain-specific neutralization or strain-specific antibody effector functions. Our study offers the first evidence that critical gB-specific non-neutralizing antibodies have different strain specific gB recognition when displayed on the surface of a cell. This panel of non-neutralizing gB-specific mAbs generally bound with greater preference to gB genotypes 2 and 4. These findings are in concordance with phylogenetic clustering of the 5 clinically significant HCMV gB genotypes 1/2/4 and 3/5 into two supergroups [29]. The lack of knowledge regarding which HCMV gB genotype or genotypes infected the individuals from which the mAbs were isolated limits the assertions we can make about how gB genotype specific antibodies may be raised by natural infection. Clinically, prevalence of HCMV gB variants may be influenced by geography, immune status, and prior infection with HCMV of other gB variants [38–40]. Future efforts might address the question of gB variant specific non-neutralizing recognition and antibody function by utilizing mAbs isolated from individuals that have been tested for endogenous viral strains or have known exposure to a specific gB genotype variant.

A notable finding from studying full-length gB-binding mAbs that may be AD3- or MPER-specific, is their breadth of membrane associated gB binding. These mAbs, 1-155, 1-237, and 3-18 demonstrate negligible gB ectodomain and postfusion gB trimer binding by both ELISA and SPR yet yield comparable signals for transfected cell binding to other gB ectodomain binding mAbs. With AD3 thought to be buried in the cytodomain [12], these findings raise the question: how do presentation of gB epitopes differ as a soluble protein versus a membrane-associated protein? As a viral fusogen, gB is thought to undergo transformation from a prefusion conformation to a postfusion conformation to facilitate entry into a host cell [21, 41]. While this study does not define a specific prefusion structure, it does highlight how a distinct prefusion-like gB on a cell surface may expose different epitopes than soluble postfusion gB, potentially accounting for improved membrane associated gB binding of AD3-specific mAbs. Notably, it was recently reported that the ability of gB/MF59 vaccine sera to bind to gB transfected cells predicts risk of HCMV acquisition [19]. Taken together, these findings warrant further investigation of gB conformation-specific antibody binding, to parse out the epitope binding specificity and effector functions of gB vaccine-elicited Abs.

This study demonstrates the first effort to identify gB domain specificity and function of antibodies that mediate Fc receptor functions like ADCP and ADCC elicited by natural HCMV infection. While little is known about gB-specific non-neutralizing functions in naturally-infected individuals, investigations of gB/MF59 vaccinees has shown this class of antibody may indeed be a desirable target of vaccines that aim to protect against primary HCMV acquisition [13]. It is now evident that non-neutralizing Abs are not limited to traditional “non-neutralizing” epitopes [12, 17]. Even more, these mAbs, while preferentially binding to certain gB genotypes, retain the ability to bind across a spectrum of gB genotype variants. Ultimately this work contributes to the field of HCMV vaccinology by emphasizing the impact of epitope binding specificity, binding strength, and genotype breadth amongst functional non-neutralizing gB-specific mAbs. By improving our knowledge of gB immunogenicity elicited by natural infection, and the specificity of antibody responses that may mediate key functions other than traditional neutralization, this work informs rational design of new HCMV vaccines that aim to reduce the devastating burden of HCMV disease in both congenital infection and transplant settings.

## Acknowledgements

The authors recognize Justin Pollara and Whitney Edwards for their assistance with the antibody dependent cellular cytotoxicity assays. The gB-specific monoclonal antibodies were kindly provided by Dai Wang at Merck through a collaboration with Zhiqiang An’s laboratory at the University of Texas Health Science Center at Houston. This work was supported by grants from Merck Research Labs and the Welch Foundation (AU-0042-20030616 to Z.A), NIH/ National Institute of Allergy and Infectious Disease R21 (R21AI136556 to S.R.P). Dr Permar consults for Merck, Pfizer, Sanofi, and Moderna vaccine programs and has sponsored programs from Merck and Moderna around CMV vaccine immunity. A patent application covering the mabs described in the article has been submitted by Merck &Co., Inc. and University of Texas. D.W. is a current employee and stockholders of Merck & Co., Inc.

